# Cortical dopamine reduces the impact of motivational biases governing automated behaviour

**DOI:** 10.1101/2021.09.09.459267

**Authors:** Vanessa Scholz, Roxanne W. Hook, Mojtaba Rostami Kandroodi, Johannes Algermissen, Konstantinos Ioannidis, David Christmas, Stephanie Valle, Trevor W. Robbins, Jon E. Grant, Samuel R. Chamberlain, Hanneke EM den Ouden

## Abstract

Motivations shape our behaviour: the promise of reward invigorates, while in the face of punishment, we hold back. Abnormalities of motivational processing are implicated in clinical disorders characterised by excessive habits and loss of top-down control, notably substance and behavioural addictions. Striatal and frontal dopamine have been hypothesised to play complementary roles in the respective generation and control of these motivational biases. However, while dopaminergic interventions have indeed been found to modulate motivational biases, these previous pharmacological studies used regionally non-selective pharmacological agents. Here, we tested the hypothesis that frontal dopamine controls the balance between Pavlovian, bias-driven automated responding and instrumentally learned action values. Specifically, we examined whether selective enhancement of cortical dopamine either (i) enables *adaptive* suppression of Pavlovian control when biases are maladaptive; or (ii) *non-specifically* modulates the degree of bias-driven automated responding. Healthy individuals (n=35) received the catechol-o-methyltransferase (COMT) inhibitor tolcapone in a randomized, double-blind, placebo-controlled cross-over design, and completed a motivational Go NoGo task known to elicit motivational biases. In support of hypothesis (ii), tolcapone globally decreased motivational bias. Specifically, tolcapone improved performance on trials where the bias was unhelpful, but impaired performance in bias-congruent conditions. These results indicate a non-selective role for cortical dopamine in the regulation of motivational processes underpinning top-down control over automated behaviour. The findings have direct relevance to understanding neurobiological mechanisms underpinning addiction and obsessive-compulsive disorders, as well as highlighting a potential trans-diagnostic novel mechanism to address such symptoms.

## Introduction

We generally feel that we are in control of our actions and make our decisions rationally. Yet, many of us eat that extra slice of cake, buy that expensive phone, or fail to save sufficiently for our retirement. While our behaviour is indeed to a large extent driven by ‘rational’ (instrumental) learning from experience, a key observation is that motivational prospects shape our behaviours in a seemingly hardwired way: the promise of rewards invigorates behaviour, while we hold back under the threat of punishment ^1–4^. These motivational biases are thought to simplify decision-making by providing sensible default actions (‘priors’) ^4^. Such decision heuristics can be particularly helpful in situations requiring rapid responding, or in an unfamiliar environment ^5^. Still, it has long been known that these Pavlovian processes shape behaviour even when the responses they prompt are maladaptive ^6,7^. In contrast, through instrumental learning of stimulus-response-outcome contingencies we can flexibly learn which actions are advantageous in any given, specific environment, which, once learnt, will lead to more optimal choices. Thus, adaptive behaviour requires a careful balance between a fast but inflexible Pavlovian ‘controller’, and an instrumental ‘controller’ that flexibly but more slowly learns adaptive behaviour in specific environments. Abnormalities in motivational processing have been implicated in clinical disorders characterised by habitual behaviours that are functionally severely impairing, for example in substance and behavioural addictions ^8,9^ as well as disorders from the obsessive-compulsive spectrum ^10–12^. Furthermore, there is evidence that Pavlovian biases governing instrumental behaviour may predict psychiatric relapse and symptom progression in certain clinical contexts ^13^ and recovery ^14^.

Influential theories and computational models posit that motivational biases arise through ventral striatal dopamine action ^2,4,15–17^, based on observations that Pavlovian cues elicit dopamine release in the ventral striatum ^18,19^. Also, in humans, dopaminergic interventions can modulate the expression of motivational bias ^3,20,21^. However, these effects are puzzling in the sense that their direction is inconsistent across studies. One cause of this seeming inconsistency may lie in the systemic nature of typical human psychopharmacological interventions (e.g. L-DOPA or psychostimulants), which typically impact both striatal and prefrontal dopamine function. Indeed, in addition to an important role of the striatum eliciting motivational bias, we posit a putative role of *frontal* dopamine in controlling these biases. In this study, we leveraged a regionally specific pharmacological intervention to ask whether and how prefrontal dopamine acts in determining the degree to which motivation biases instrumental behaviour.

While most dopaminergic agents affect sub-cortical and cortical dopamine, the catechol-o-methyl transferase (COMT) inhibitor tolcapone has a highly specific effect in modulating frontal dopamine ^22^. In contrast to the striatum, where dopamine metabolism is dominated by action of the dopamine transporters, dopamine metabolism in the prefrontal cortex (PFC) primarily relies on COMT ^22,23^. The COMT enzyme plays a cardinal role in the regulation of cortical dopamine^22,23^. Evidence from pre-clinical models demonstrates that COMT knock-out leads to substantial increase in prefrontal dopamine levels in the absence of marked effects on striatal dopamine ^23,24^. Tolcapone prevents the COMT enzyme from breaking down dopamine in the PFC, leading to elevated frontal DA measured using microdialysis in rats ^25^. Tolcapone modulates aspects of flexible responding and executive control in pre-clinical and human experimental models ^25^. There is also emerging evidence that tolcapone may constitute a new therapeutic direction for disorders characterised by loss of control over habitual patterns of behaviour ^26–29^. For example, in an open-label study, over the course of 12-weeks tolcapone was associated with symptom reduction in gambling disorder, the extent of which correlated with enhancement of frontal lobe activation during an executive planning task ^28^. In a recent controlled study, two-week treatment with tolcapone led to significant improvements in OCD versus placebo ^29^. Furthermore, single-dose tolcapone has also been found to modulate activation of the right inferior frontal gyrus in people with disordered gambling, versus placebo ^26^ – a key region heavily implicated in exerting top-down control over learnt behaviours ^30–32^.

Given the selective effects of tolcapone on cortical as opposed to striatal dopamine, as well as the initial evidence indicating tolcapone may offer therapeutic promise in the treatment of disorders associated with excessive habitual patterns of behaviour, we used a single-dose challenge in conjunction with an established probabilistic reinforcement learning task. In this task, participants need to learn to make (Go) or withhold (NoGo) responding in order to obtain desired outcomes. Cues signal both the action requirement (Go / NoGo response) and outcome valence (i.e. whether for this cue a reward can be won, or rather a punishment needs to be avoided). Participants perform better for cues that require actions congruent with the outcome valence (i.e. make a Go response to win a reward, or a NoGo response to avoid a punishment) relative to incongruent cues (NoGo to win a reward, Go to avoid a punishment). This difference in performance on action-valence congruent relative to incongruent cues reflects the strength of the (ability to control the) motivational bias that prompts actions based on the cue valence. This task thus robustly evokes motivational biasing of action, which needs to be suppressed on so-called ‘incongruent’ trials to perform well. We used this motivational Go-NoGo task to characterise the role of cortical dopamine in determining the balance between automated and controlled responding in healthy volunteers.

Using a double-blind, randomized, cross-over, within-subject design, we examined whether tolcapone would facilitate a shift from bias-dictated automated behaviour towards more flexible responding, through elevation of frontal dopamine levels. Specifically, we tested the following two competing accounts. **Hypothesis 1:** Dopamine enhances suppression of Pavlovian biases when these conflict with instrumental requirements. This hypothesis follows from previous work indicating that i) the frontal cortical EEG activity predicts adaptive suppression of motivational biases within ^33^ and across ^34^ individuals, and ii) higher frontal dopamine, either through pharmacological intervention ^35^ or owing to a genetic phenotype impacting the COMT enzyme ^36^, can lead to the employment of more adaptive decision strategies. **Hypothesis 2:** Dopamine enables general disengagement from the automatic response systems, i.e. irrespective of whether biases are conducive to or interfering with selecting the correct instrumental response.

Automatic response tendencies can be suppressed by prefrontal circuits, notably the inferior frontal gyrus (IFG) ^31,37,38^, interfering with subcortical action selection processes. The IFG projects to the subthalamic nucleus (STN) and is believed to raise the threshold needed to elicit a motor response, which prevents impulsive responses ^15,39–42^. Administration of catecholamine agonists such as methylphenidate, modafinil, and atomoxetine have been found to improve response inhibition, e.g., in the stop-signal task ^43–46^. Tonic increases in IFG activation by tolcapone could thus diminish the impact of automatic, bias-driven responses and facilitate the enactment of controlled, instrumental responses^44^. Based on this literature, our second hypothesis was that tolcapone might enhance prefrontally driven response inhibition, leading to a global shift away from automatic, bias-driven responding, irrespective of whether this supports or hinders adaptive decision-making.

## Materials and Methods

### Sample

Forty-four healthy subjects meeting inclusion criteria (for an outline see Suppl. Material) took part in a double-blind, randomized, within-subjects, placebo-controlled study examining effects of a single dose of tolcapone (200 mg, dose based on previous work ^47–49^). They were recruited at two test sites, University of Cambridge (*N* = 23) and University of Chicago (*N* = 21). Additional data exclusion (see data availability), left an available sample of *N* = 35 for subsequent analysis (see Table 1).

**Table 1.**
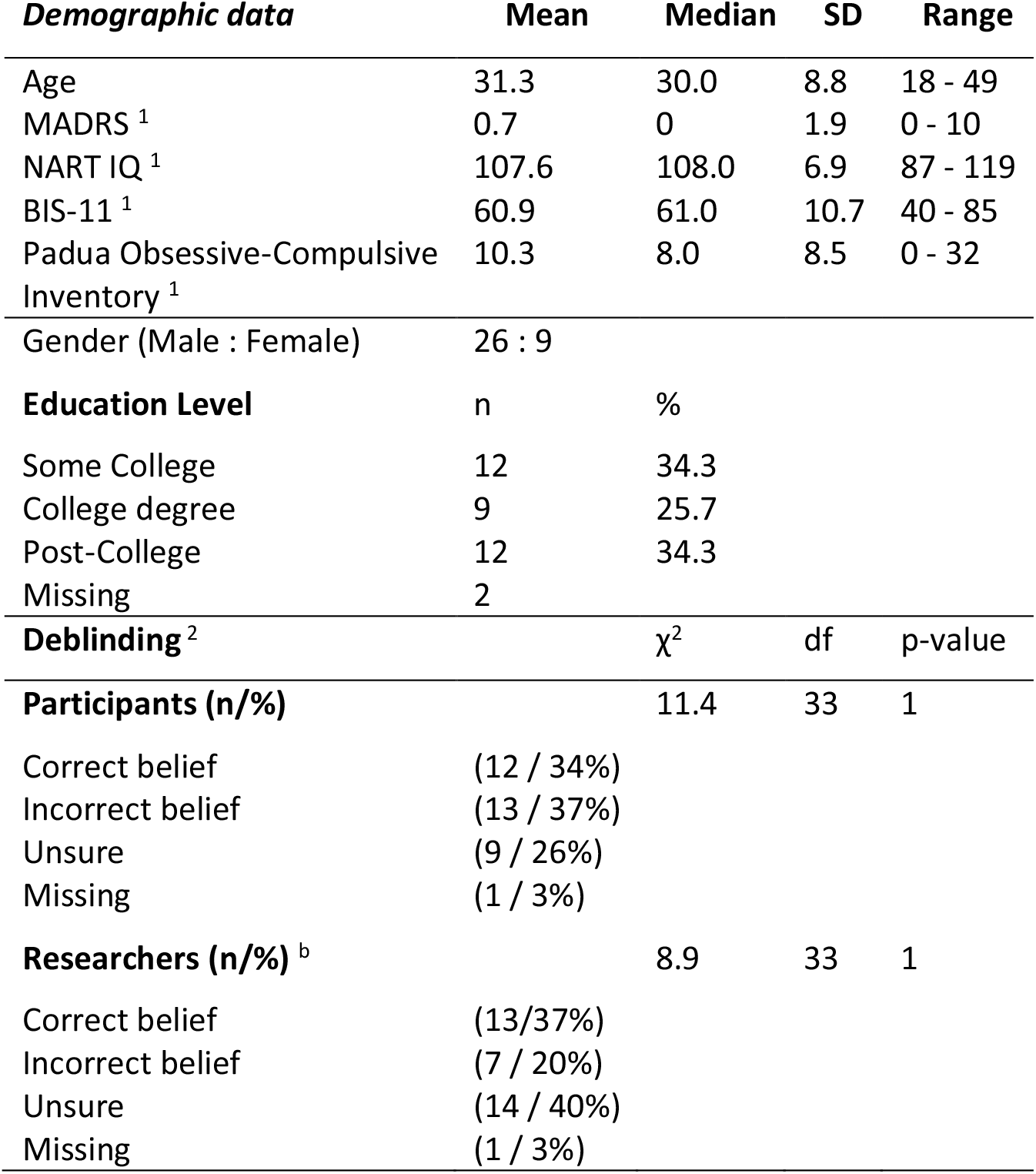
Sample characteristics and deblinding information. ^1^ scores measured at baseline testing day.^2^ After having completed the study, participants were asked to indicate their belief about when they had received the active medication, i.e. on the first or second visit; similarly, the research team was asked whether they felt a particular individual had received active treatment on the first or second visit. MADRS: Montgomery-Asberg Depression Rating Scale. NART IQ: National Adult Reading Test Intelligence Quotient; BIS-11: Barratt Impulsivity Scale 11. These variables were collected to characterize the sample in terms of IQ, and traits of impulsivity/compulsivity. χ^2^ = Pearson’s Chi-squared test

| **Demographic data** | **Mean** | **Median** | **SD** | **Range** |
| --- | --- | --- | --- | --- |
| Age | 31.3 | 30.0 | 8.8 | 18 - 49 |
| MADRS <sup>1</sup> | 0.7 | 0 | 1.9 | 0 - 10 |
| NART IQ <sup>1</sup> | 107.6 | 108.0 | 6.9 | 87 - 119 |
| BIS-11 <sup>1</sup> | 60.9 | 61.0 | 10.7 | 40 - 85 |
| Padua Obsessive-Compulsive Inventory <sup>1</sup> | 10.3 | 8.0 | 8.5 | 0 - 32 |
| Gender (Male : Female) | 26 : 9 |  |  |  |
| **Education Level** | **n** | **%** |  |  |
| Some College | 12 | 34.3 |  |  |
| College degree | 9 | 25.7 |  |  |
| Post-College | 12 | 34.3 |  |  |
| Missing | 2 |  |  |  |
| **Deblinding <sup>2</sup>** |  | **$\chi^2$** | **df** | **p-value** |
| **Participants (n/%)** |  | 11.4 | 33 | 1 |
| Correct belief | (12 / 34%) |  |  |  |
| Incorrect belief | (13 / 37%) |  |  |  |
| Unsure | (9 / 26%) |  |  |  |
| Missing | (1 / 3%) |  |  |  |
| **Researchers (n/%) <sup>b</sup>** |  | 8.9 | 33 | 1 |
| Correct belief | (13/37%) |  |  |  |
| Incorrect belief | (7 / 20%) |  |  |  |
| Unsure | (14 / 40%) |  |  |  |
| Missing | (1 / 3%) |  |  |  |

### Experimental procedure

Participation consisted of two test days separated by a period of at least one week in-between test sessions to ensure full drug washout. The first test day included a clinical interview, a medical screening, and clinical questionnaires (outlined in more detail in the suppl. material). Participants then orally received a capsule containing either 200 mg of tolcapone or a placebo. Capsules were manufactured by an independent pharmacy and were of identical appearance and weight; the randomization was done using a computer-generated randomization algorithm by the independent pharmacy. Peak plasma levels of tolcapone are achieved approximately one hour post administration and its half-life is around 4 hours ^50^. After one hour, subjects performed the motivational Go NoGo task. This was conducted as part of a broader study also including neuroimaging, results for which will be reported separately. After completion of the study, participants and experimenters were debriefed about what session they believed comprised the active treatment, enabling us to assess actual success of the blinding procedure. We confirmed successful blinding, i.e. individuals’ ability to indicate the session of active treatment did not differ from chance, for participants and experimenters (see Table 1). Participants were reimbursed with £75/100$ for study completion, plus additional travel expenses. Before taking part, all participants provided informed consent. Both ethics committees approved the study procedure (East of England- Cambridge East Research Ethics Committee IRB: 16/EE/0260 and Ethics Committee University of Chicago, IRB 16-0738), which was in accordance with the Declaration of Helsinki 1975.

### Motivational Go NoGo task

We employed a well-established reinforcement learning task to evoke and measure motivational biases (identical to van Nuland et al. (2020) ^21^, originally adapted from Guitart-Masip et al. (2011) ^51^. In the motivational Go NoGo task, participants were presented with one cue out of four possible cue categories (Go2Win, Go2Avoid, NoGo2Win, NoGo2Avoid), on each trial (1300 ms) and needed to decide on a Go (button press) or a NoGo response (withholding a button press) (Figure 1) before cue offset. On each test day, the task consisted of two blocks of 80 trials with each cue category presented 40 times, thus 160 trials in total per test day. For each test day, a different cue set was employed to alleviate training effects. On the first day, participants performed practice trials before the task.

**Figure 1:**
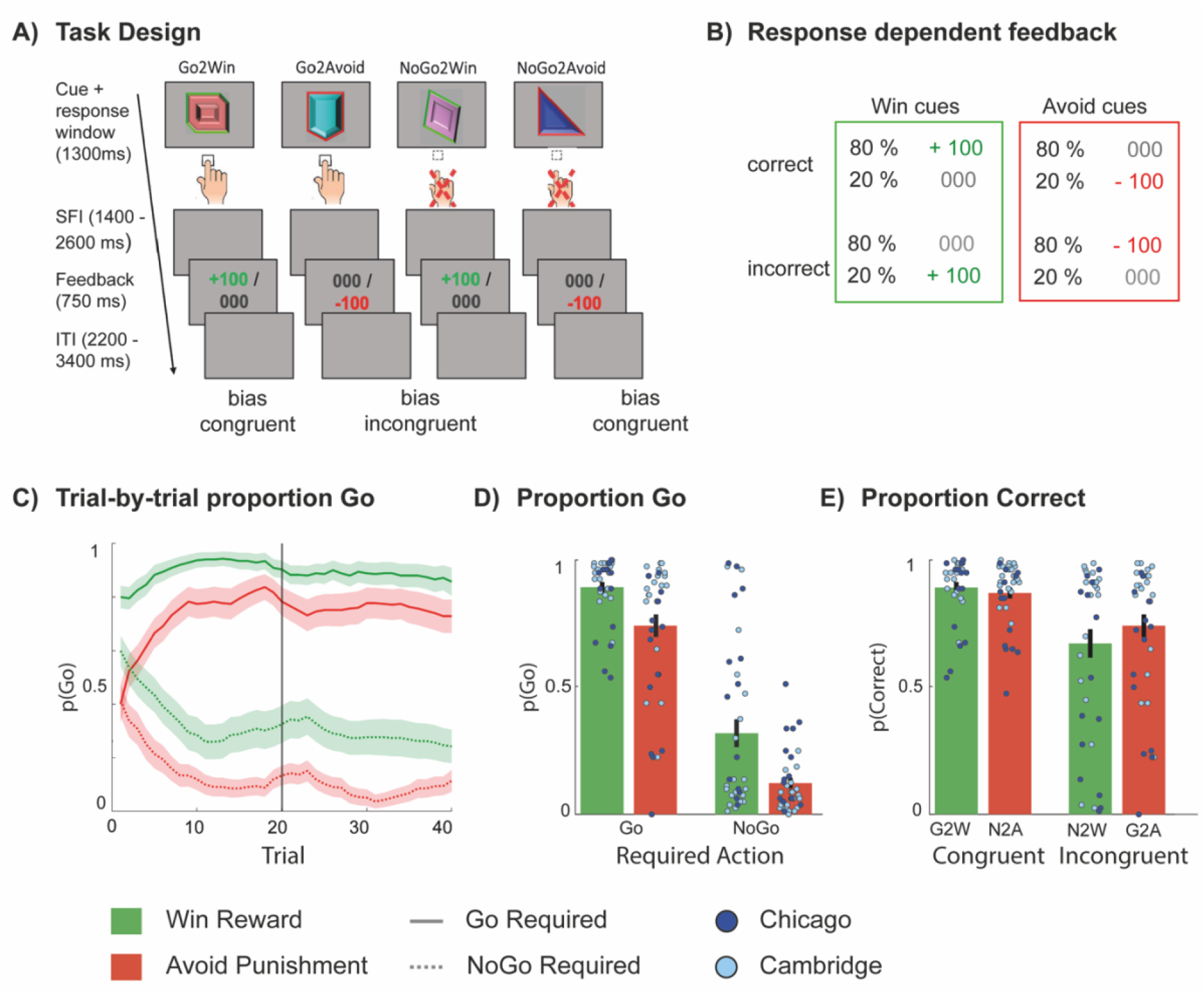
Motivational Go-NoGo task design and overview of main task effects. **A)** Go NoGo task trial sequence for each of the four cue categories: Go-to-Win, Go-to-Avoid, NoGo-to-Win, and NoGo-to-Avoid. Go-to-Win and NoGo-to-Avoid are bias congruent cue categories, as their action requirement is in line with the stimulus-response coupling strengthened by the motivational bias. Go-to-Avoid and NoGo-to-Win are bias-incongruent response-stimulus couplings, which are usually harder to execute for participants. On each trial, a cue was presented for 1300 milliseconds (ms) and subjects could decide to make a Go response by pressing a button or choosing a NoGo response by withholding a response. After this, subjects were presented with the outcome (reward, neutral, punishment) for 750 ms, the valence of which was determined by the cue category and the probabilistic feedback schedule. The inter-trial-interval (ITI) was 2200 - 3400 ms, in steps of 200 ms **B)** The feedback contingencies for this task version were 80% : 20%. **C)** Trial-by-trial behaviour. Depiction of the probability of making a Go response, P(Go), (± SEM) and plotted with a sliding window of 5 trials for Go cues (solid lines) and NoGo cues (dashed lines) across trials per cue category, here collapsed across both treatments (tolcapone and placebo). Choice biases are evident from the first trial onwards, as the green lines characterizing P(Go) for Win cues are always above the red lines depicting the probability of making a go response for cues requiring a NoGo response as optimal action choice. **D)** Probability of making a Go response for each cue condition, grouped by required action. Learning is evident from the increased proportion of ‘Go’ responses to Go cues. Motivational/ Pavlovian biases is evident from the reduced probability of Go responses to Avoid cues **E)** Probability of making a correct response (i.e. 1-pGo for NoGo cues), reorganised so that now bias-congruent and bias incongruent cues are grouped together. Note that this means that the data plotted here are the same as in panel D for Go cues, and the inverse for NoGo cues. This more clearly illustrates the reduced accuracy on bias-incongruent cues, regardless of action requirement. Cue categories abbreviated as follows: G2W = Go to Win, G2A = Go to Avoid Punishment, N2W = NoGo to Win, N2A = NoGo to Avoid Punishment

For Win cues, participants could receive a reward (desired) or neutral feedback (non-desired). In contrast, for Avoid Punishment cues, participants could receive either a neutral feedback (desired) or a punishment (non-desired). Outcome valence was signalled by the colour of the cue edge (red for Avoid cues, green for Win cues). Feedback was displayed in the centre of the screen for 750 ms. (Figure 1). Guided by this feedback, participants had to learn by trial and error which response was best for each cue. Feedback was probabilistic: A correct response (e.g. a Go response for a Go2Win or Go2Avoid cue) resulted in the desired outcome on 80% of trials, while for 20% of correct responses, participants received a non-desired outcome. Vice versa, incorrect responses led to the non-desired outcome on 80% of trials, and to the desired outcome on the remaining 20% of trials. Importantly, Go2Win and NoGo2Avoid cues are bias-congruent cues, as their required action is in line with the actions prompted by the valence of the cue (i.e. motivational bias). Accuracy on these congruent trials is expected to be high. In contrast, Go2Avoid and NoGo2Win cues are bias-incongruent cues, i.e. their instrumental action requirement conflict with the action facilitated by motivational biases resulting in reduced accuracy (Figure 1D/E and Suppl.).

Trials were interspersed with inter-trial-intervals (ITI) 2200 - 3400 ms, in steps of 200 ms. Each step size was presented the same number of times, for each cue and step-size. Within each cue, the temporal sequence of ITIs was randomised. Cue-feedback intervals were also jittered using the same procedure, now using a range of 1400-2600 ms, again with stepsize of 200 ms.

### Data availability

Data inspection and analyses were conducted by team members who remained blind to drug condition, until after all analyses were completed. We analyzed participants who completed both sessions (placebo and tolcapone) and for whom sufficient behavioural data were available (N = 35). From the original sample of 44 subjects: 5 participants had to be excluded due to technical issues resulting in data loss, 2 had been accidentally presented with the same cue set twice rendering their performance incomparable to other participants and 1 participant did not return for the second visit. Data were subsequently screened in terms of data quality and missing data (see Suppl. for the *a-priori* defined criteria). Based on this assessment, 1 participant was excluded due to missing data.

### Analytic Approaches

We used two complementary statistical approaches to analyse the data. The first used conventional logistic mixed-effects models; and the second tested computational models based on a priori literature.

First, we analyzed how the probability of making a Go response P(Go) was affected by the following three within-subject factors and their interactions: required action (Go, NoGo), valence (Win; Avoid), and treatment (tolcapone, placebo). We focused on the following effects of interest: i) Main effect of required action. This reflects a differential tendency to make a Go response as a function of the required (Go or NoGo) response, capturing learning to make the correct response. ii) Main effect of valence. This reflects a differential tendency to make a Go response to Win vs. Avoid cues, capturing motivational bias. iii) Valence x Drug interaction. This reflects a differential motivational bias as function of tolcapone administration. As data was acquired at two sites, we included a between subject factor ‘Site’, as a control variable, which was allowed to interact with all model terms of the initial model (see supplemental materials for the full model equations. Next, in a follow-up analysis, we also tested whether the (effect of tolcapone on) motivational bias was constant over time, by adding ‘task block’ as a within-subject factor interacting with the above effects.

Finally, we verified that testing order (tolcapone vs. placebo on session 1) did not interact with the observed Valence x Drug interaction, by including between-subject Testing Order (refer to Suppl. for full report of results). For general interest, we also report analyses of reaction time (RT) data (see Suppl.). All models contained the full random effects structure for the within-subject variables. Generalized logistic mixed-models analysis was conducted using lme4, version 1.1-23 ^52^ in R 4.0.2. Statistical significance was determined as p-values with α< 0.05, two-sided.

Second, to dissect the computational mechanisms sub-serving motivational action bias and evolving instrumental learning, we fitted three hierarchically nested reinforcement learning (RL) models ^3^. Model equations are provided in the Supplements. In brief, M1 was a basic Rescorla Wagner model^53^ and contained a parameter for feedback sensitivity and a learning rate that together would learn the Q values. M2 extended model 2 with a ‘Go bias’ parameter *b* that captured the overall tendency to make Go responses. M3 extended M2 with a motivational bias parameter π which could capture the differential tendency to make more Go responses to Win cues. We then established through model comparison (Suppl.), whether additional model parameters increased model evidence. After establishing the winning model (M3), we extended this winning model to model M4, where all parameters were allowed to be modulated by tolcapone. Model M4 comprised two separate parameters sets for the placebo (ρ_pla_, ε_pla_, b_pla_, π_pla_) and drug session (ρ_tolc_, ε_tolc_, b_tolc_, π_tolc_). We then ran a second model comparison comparing models M1-4 to establish evidence for tolcapone modulating the model parameters. To assess the specific effect of tolcapone on each model parameters, we compared parameters of both drug conditions while controlling for site.

## Results

### Generalized linear mixed models for choice data

We regressed participants’ choices onto cue valence, required action, and drug condition, with test site as between-subjects factor. We observed significant main effects of required action, indicating that participants learned the task; and valence, indicating that participants’ choices were affected by motivational biases, with more Go responses to Win cues than Avoid cues (Table 2). The interaction of required action and valence was non-significant, providing no evidence for motivational biases differing in size for Go vs. NoGo cues.

**Table 2.** Full statistics report of the main mixed-effects regression model for choice data, a effects analysis to characterize the treatment effect. Abbreviations: SE = Standard Error

| | $\beta$ estimates | SE | $\chi^2$ | p-value |
| --- | --- | --- | --- | --- |
| **Main effects** |  |  |  |  |
| valence | -0.801 | 0.15 | 27.4 | <.001 *** |
| required action | 1.954 | 0.20 | 95.3 | <.001 *** |
| drug | -0.091 | 0.08 | 1.4 | .2 |
| site | -0.006 | 0.12 | <0.01 | 1 |
| **Interaction effects** |  |  |  |  |
| required action x valence | -0.010 | 0.07 | 0.02 | 0.9 |
| drug x site | 0.061 | 0.08 | 0.6 | 0.4 |
| valence x drug | -0.200 | 0.08 | 6.1 | 0.01 * |
| valence x site | 0.139 | 0.14 | 0.8 | 0.4 |
| required action x drug | 0.102 | 0.12 | 0.8 | 0.4 |
| required action x site | 0.570 | 0.20 | 8.1 | 0.004 ** |
| valence x drug x site | 0.070 | 0.08 | 0.8 | 0.4 |
| required action x drug x site | -0.991 | 0.12 | 0.7 | 0.4 |
| valence x required action x drug | 0.032 | 0.06 | 0.3 | 0.6 |
| valence x required action x site | 0.039 | 0.07 | 0.3 | 0.6 |
| valence x req. action x drug site | 0.007 | 0.06 | 0.01 | 0.9 |
| **Post hoc simple effects (Win – Avoid)** |  |  |  |  |
| Placebo | -0.954 | 0.19 | 26.4 | <.001 *** |
| Tolcapone | -0.583 | 0.14 | 14.7 | <.001 *** |

There was a significant Drug x Valence interaction effect (χ^2^(1) = 6.1, p-value = .01) indicating that the main modulatory effect of cue valence on ‘Go’ responding, i.e. the motivational bias, was modulated by tolcapone. The direction of this effect was such that under tolcapone, there was less bias than under placebo (c.f. post-hoc simple effects). Importantly, there was no ‘Required Action x Valence x Drug’ interaction (χ^2^(1) = 0.3, p-value = .6). This meant that there was no evidence for the degree of biased responding to be different as a function of required action (i.e. the degree of ‘Go’ responding for Win cues increased regardless of whether a Go was required or not, i.e. whether the bias was congruent or incongruent with the action requirements. Thus, these results support Hypothesis 2 that tolcapone globally reduced motivational bias. Further examining the effect of task block, this effect of tolcapone on motivational bias was not constant across the task (Block x Valence x Drug: χ^2^(1) = 4.6, p-value = .03) (report of full analysis results in Suppl.). Post-hoc simple effects on each block showed that motivational biases were significantly reduced under tolcapone in the first block only (Block 1: χ^2^(1) = 7.6, p-value = 0.006; Block 2: χ^2^(1) < 0.01, p-value = 1 c.f. Figure 2 E-F).

**Figure 2.**
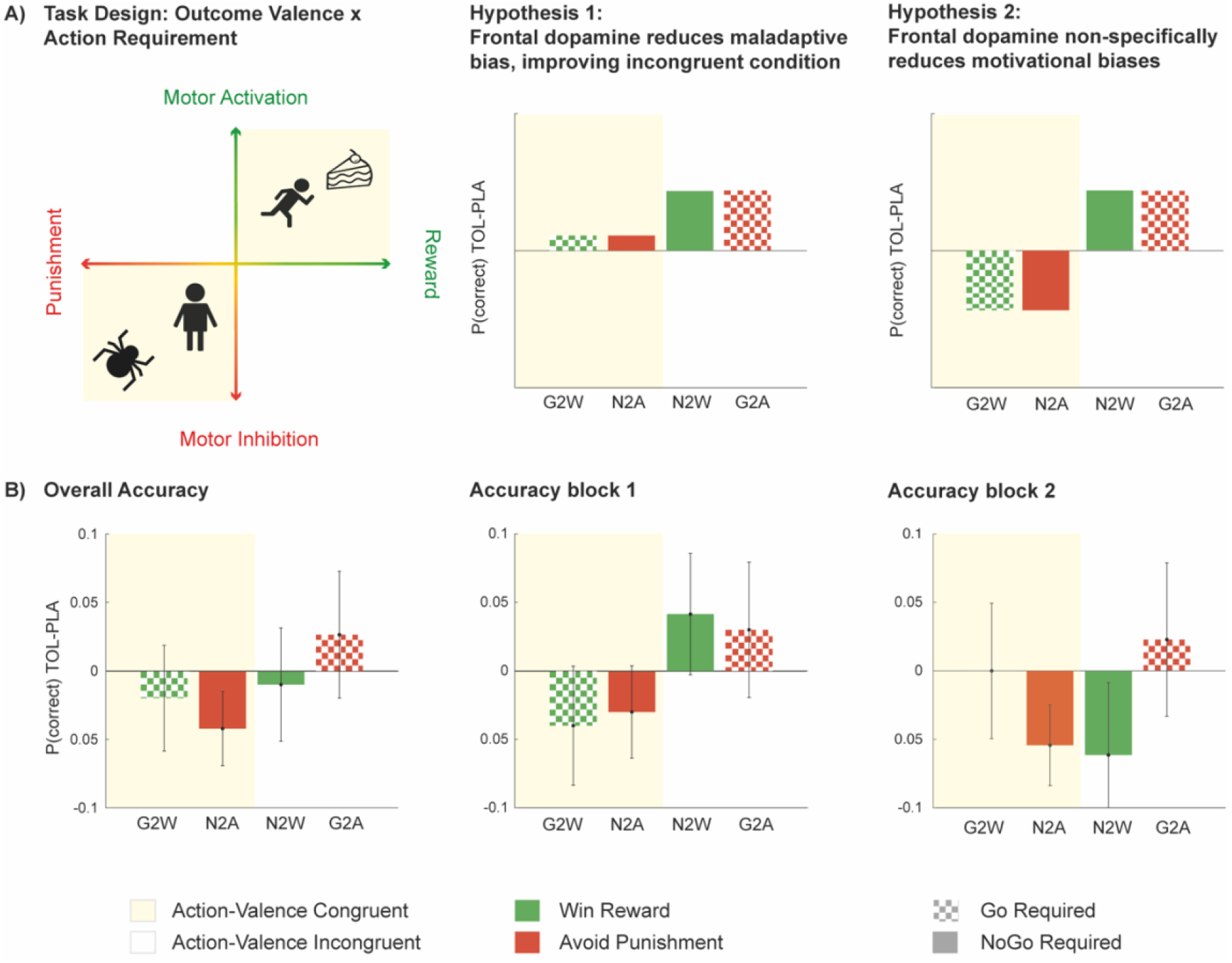
Hypothesised and measured effect of tolcapone administration: **A)** Illustration of task design to capture motivational biases- through coupling of the orthogonalized axes of motivational valence (Reward, Punishment) and action (motor activation | Go) or (motor inhibition | NoGo). Yellow: valence-action bias congruent responses is required; White: bias-incongruent responses is required **B+C)** Predicted change in choice accuracy following tolcapone administration relative to placebo, for each of the 4 conditions. The right 2 panels represent the hypothesised effects of tolcapone. **Hypothesis 1**: Tolcapone enhances adaptive control, i.e. suppresses Pavlovian bias on incongruent trials, thereby increases the proportion of correct responses (accuracy) on incongruent trials. Speculatively, performance on congruent trials may improve also. **Hypothesis 2**: Tolcapone promotes a general shift away from automated responding, reducing bias overall. This would lead to improved choice accuracy on incongruent trials (as for Hypothesis 1), but crucially, to reduced choice accuracy for congruent trials (highlighted in yellow). **B)** Data: Mean (±SED) accuracy, i.e. proportion of correct responses, under tolcapone relative to placebo, shown across all trials, for the first half of the trials (block 1) only, and for the 2^nd^ half of the trials (block 2) only. In line with hypothesis 2, performance on congruent trials is reduced, while performance on incongruent trials is reduced. This is particularly evident for block 1. *G2W = Go to Win, G2A = Go to Avoid Punishment, NG2W = NoGo to Win, NG2A = NoGo to Avoid Punishment*

### Computational modelling and model comparison

Replicating many previous studies ^3,51,54^, base model comparison (M1-M3) indicated the highest evidence for model M3, which extended a basic reinforcement learning model with ‘go’ and motivational bias parameter (Figure 3; model frequency: 42.9%; protected exceedance probability (PXP) =0.7). Addition to the model space of an extension of this winning model with separate tolcapone and placebo parameters provided very strong evidence that this again improved the model (M4 model frequency = 60.5 %, PXP = 1.0, see also Suppl. Table S4)

**Figure 3.**
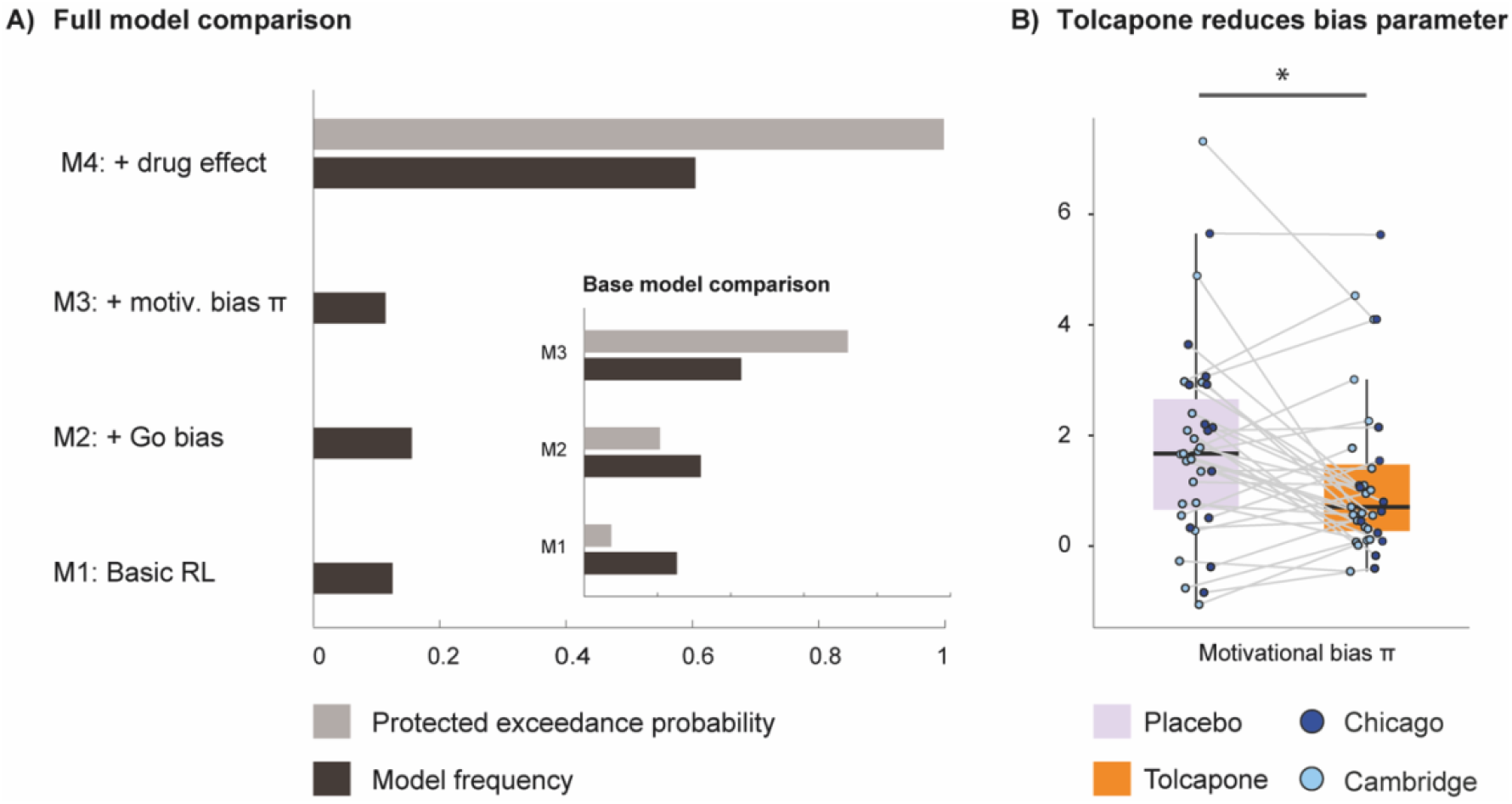
Tolcapone induced changes in model parameter estimates. A) Full model comparison showing Model M4 including four parameters, namely feedback sensitivity, a learning rate, a Go bias and a motivational bias parameter to outperform the other three base models. As a small inset, the base model comparison is shown. Here, model M3 outperformed the simpler models M1 and M2. Model frequency and protected exceedance probability were employed as model fit indices. B) The π parameter capturing effects of motivational biases was significantly reduced under tolcapone administration. The remaining parameters feedback sensitivity, learning rate and Go bias were not significantly affected by tolcapone, indicated by p-values > .01 for all main effects of condition or interaction terms.

The motivational bias parameter π was significantly reduced under tolcapone relative to placebo (χ^2^(1) = 5.4, p-value = .02; Figure 3) and this effect did not differ as a function of test site (Drug x Testing site: χ^2^(1) < 0.1, p-value =.9; for full report for the interaction of the other parameters with testing site, see Suppl. Table S5). We also verified that there were no significant tolcapone-induced differences for any of the other parameters (all p-values > .1, see Suppl. Fig S3 and Table S5). Finally, through data simulation using the winning model’s estimated parameters, and refitting them to the simulated data, we were also able to recover the tolcapone effect on the bias parameter π (χ^2^(1) = 6.3, p-value = .01 (see supplemental for more information on absolute model fit and effect recovery).

## Discussion

This study’s primary goal was to examine the impact of a cortical dopamine challenge on motivational biases using the COMT inhibitor tolcapone, to evaluate two alternative hypotheses regarding the role of cortical dopamine in motivational processing. The first hypothesis posited adaptive bias reduction under tolcapone, supressing motivational biases whenever instrumental and Pavlovian control conflicted, while the second hypothesis proposed a global reduction in motivational biases, regardless of whether these aligned with or opposed instrumentally learnt action values. Our key finding was that tolcapone significantly decreased motivational biases across both bias-congruent and incongruent Pavlovian-instrumental trials, supporting the second hypothesis that cortical dopamine non-selectively dampens the impact of motivational biases on behaviour. This effect was established using both conventional statistical analysis and computational modelling. Due to the global bias reduction, tolcapone did not generally improve performance, but rather decreased performance on bias-congruent trials, while improving performance on bias-incongruent trials. We objectively confirmed that the study was successfully double-blinded.

Our findings accord with previous findings on catecholaminergic agonists improving response inhibition ^11,41,55^. A stronger tonic drive from IFG via the subthalamic nucleus might raise response thresholds in the striatum and in this way prevent the enactment of automatic, prepotent responses ^31,37,39,40,56^. Importantly, modulation of frontal dopamine can thus have opponent effects to modulation of striatal dopamine. A recent study in rodents directly compared effects of dopamine transporter (DAT) blockade, with DAT putatively forming the primary mechanism of striatal dopamine clearance, with COMT inhibition. In this study DAT blockade selectively impaired, and COMT inhibition improved performance after reward reversals ^57^. This finding is particularly noteworthy given the opposite effects of two interventions that both increase dopamine, yet presumably in different locations, namely the striatum and prefrontal cortex respectively. Stimulation of the meso-cortical dopamine pathway in this study also provides a clue as to the kind of cognitive effects we may expect to see from COMT inhibition both in this study as well as in the clinical domain. Especially noteworthy is that COMT inhibition did not affect fast ventral-tegmental-evoked dopamine transients in the PFC^57^, despite well-known associations between COMT activity and dopamine levels recorded over longer timescales by microdialysis ^25^ The observation that COMT inhibition may affect dopamine on longer rather than shorter timescales can provide a biological level understanding of the observation in the current study that COMT inhibition through tolcapone affected the *overall* tendency of biased responding, rather than fast, trial-specific adaptive modulation. This effect could be of particular clinical relevance for disorders characterized by an excessive reliance of automated and habitual responding, as this generalized effect might not only modulate biased responding as reported here, but potentially also affect reliance on habits, i.e. reduce over-habitual behaviour.

Importantly, the effect of tolcapone was present only in the first task block on each study visit, when instrumental learning had not yet reached asymptote (c.f. Figure 1C). This is relevant because Pavlovian biases have been shown to affect behaviour most strongly when there is high uncertainty about the instrumentally learnt action values^58^, in line with more general ideas that the balance between decision controllers is determined by their relative (un)certainty ^58–60^. As such, during the early stages of the task, individuals are more prone to rely on default priors, i.e. motivational action biases, which have been established through experience. However, as we repeatedly observe the consequences of our actions in the current task environment, the instrumental controller ‘gains confidence’ in the learnt action values associated with each cue, and takes over as the dominant system guiding choice. Here, we then show that boosting frontal dopamine causes individuals to reduce this early reliance on the Pavlovian system. This earlier shift could be due to perceived increase of control, or perceived down-weighting of the cost of reliance on a more cognitively effortful strategy ^61–63^. Support for this also comes from a study by Westbrook and colleagues (2020), who showed changes in striatal dopamine to promote the willingness to exert cognitive effort on a cognitive task by altering the subjective cost-benefit ratio of cognitive control in favor of benefits ^64^.

An alternative interpretation of our findings of tolcapone-induced bias reduction is that tolcapone reduces the integration of Pavlovian and instrumental knowledge. Neurally, this integration could be implemented through interaction between the orbitofrontal cortex (OFC), processing Pavlovian values, and the rostral anterior cingulate cortex, processing instrumental action values. This idea is supported by a recent study in marmoset monkeys by Duan et al (2021)^65^ showing that the rostral anterior cingulate cortex is necessary for detecting instrumental control of actions over outcomes, while the anterior orbitofrontal cortex (OFC) mediates Pavlovian influences on goal-directed behaviour. In line with this we have also recently shown that BOLD activity in the orbitofrontal / ventromedial prefrontal cortex predicts the degree of valence-induced invigoration ^66^. This notion would align with previous work showing that modulating frontal dopamine can reconfigure connectivity patterns between OFC and other brain regions suggesting a key role in shaping functional brain circuitry ^67^. More specifically in relation to the function of prefrontal COMT activity, COMT genetic phenotype modulated functional connectivity patterns of frontal regions including the anterior cingulate cortex with higher enzymatic activity corresponding to stronger connectivity compared to lower COMT activity^68^. Future studies should investigate whether the reported changes in biased responding under tolcapone corresponds to changes in functional connectivity strength during the task.

Altered motivational biases have been linked to psychiatric disorders such as substance and behavioural addictions ^8,9^ as well as obsessive-compulsive related disorders ^10–12^. Given the observed effects of tolcapone on motivational processing in healthy volunteers, it may be a valuable avenue for future work to examine effects of tolcapone on motivational processing and symptoms in psychiatric conditions characterised by over-expression of automated behaviours. In addition to tolcapone, other brain-penetrant COMT inhibitors are likely to become available in future ^69^. The clinical potential of COMT inhibitors is suggested by recently reported improvements following two-week tolcapone treatment in OCD, as compared to placebo; as well as by other contextual studies in healthy controls suggestive of cortically-relevant cognitive effects ^28,29,70^.

Whilst we show robust effects of tolcapone, there are some limitations that need to be considered. First, this was a single-dose study in healthy volunteers; as such, findings may differ if smaller/larger pill doses are used, or medication is administered over a different time frame; or may also vary as a function of basal levels of cortical dopamine. Indeed, pharmacological dopaminergic effects on the trade-off between cognitive flexibility and stability have often been shown to depend on baseline dopamine levels such that dopamine levels and performance on set-shifting and reversal tasks followed an inverted U-shape ^56,71–74^. Therefore, future work may wish to include larger number of sites and sample sizes to identify variables that may contribute to differential effects of tolcapone across individuals.

In sum, we showed that tolcapone significantly reduced the reliance on automatic behaviour in healthy individuals, in an experimental medicine study using a laboratory-based task assessing motivational processes. The data suggest that cortical dopamine enhancement using COMT inhibitors merits further research as a candidate trans-diagnostic treatment approach for disorders characterized by excessive habits. Employing computational modelling to characterize the latent mechanism underlying dopamine induced changes in motivational choice behaviour under tolcapone, this study helps to address a previous translational gap. Future work should use similar approaches alongside clinical outcome measures to confirm mechanisms in clinical contexts using tolcapone and other COMT inhibitors.

## Acknowledgements

This research was funded by Wellcome via grants [110049/Z/15/Z & 110049/Z/15/A] to SRC. For the purpose of open access, the author has applied a CC BY public copyright licence to any Author Accepted Manuscript version arising from this submission. VS was supported by a research scholarship by the German Research Foundation (SCHO 1815/1-1 & SCHO 1815/2-1). HEMdO was supported by a Vidi Award from the Netherlands Organization for Scientific Research (175.450). The funders had no role in the study design, data collection and analysis, decision to publish or preparation of the manuscript.

## Authors Contribution

SRC was the Chief Investigator on this study, responsible for the over-arching protocol design and the conduct of the trial in the UK. JEG was Principal Investigator at the University of Chicago and responsible for the conduct of the trial in the USA. HEMdO designed the cognitive paradigm. HEMdO, KI, JEG, TWR, DC and SRC all contributed to aspects of the study design. RWH and SV collected the data. VS and HEMdO designed and conducted data analysis. VS; JA, MRK, JEG, SRC and HEMdO discussed statistical analyses and results. VS, SRC, JA, HEMdO wrote the manuscript. All authors commented on or edited the manuscript.

## Disclosure

SRC receives an honorarium for editorial work at Elsevier; and previously consulted for Promentis. JEG has received research grants from the TLC Foundation for Body-Focused Repetitive Behaviors, and Otsuka, Biohaven, and Avanir Pharmaceuticals. TWR discloses consultancies with Cambridge Cognition, Arcadia, Greenfield Bioventures, Heptares, Takeda, Lundbeck, Merck, Sharp and Dohme. Royalties with Cambridge Cognition. Research Grants with Shionogi and GlaxoSmithKline and editorial honoraria with Springer Nature and Elsevier.The remaining authors report no disclosures of relevance.

## Competing Interests

The authors declare no competing interests.

