## Supplemental material for "Cortical dopamine reduces the impact of motivational biases governing automated behaviour"

### Supplemental Materials - Table of Contents

### Sample Description

#### Sample and site description

Individuals were considered eligible if they i) were between 18-49 years of age, ii) had no prior history of neurological disorders (e.g. Parkinson's Disease) iii) had no medical conditions or medication considered incompatible with drug administration, iv) had no current, clinically relevant depression (Montgomery-Asberg Depression Rating Score: MADRS >20) (Montgomery & Asberg, 1979), had no history of bipolar disorder, psychotic disorder or a diagnosis of Borderline Personality Disorder, vi) reported no recent use of illicit substances (self-report and as confirmed by urine screen) and fulfilled MRI inclusion criteria.

#### Clinical assessment, neuropsychological testing and clinical questionnaires

Clinical assessment was carried out by a trained rater and comprised a structured assessment of medical and psychiatric history, including medications, the National Adult Reading Test to quantify IQ [1], and the Montgomery Asberg Depression Rating Scale (MADRS) to quantify depressive symptoms [2]. Clinical interview included the Mini International Neuropsychiatric Inventory (MINI) to identify mental disorders[3], which were exclusionary. Participants then completed the Barratt Impulsiveness Scale (BIS-11) [4] and the Padua Inventory (PI-WSUR) [5] to characterize their self-reported levels of impulsivity and compulsivity. A characterization of the sample including the demographic data and questionnaire scores can be found in table S1.

| **Table S1** | **Cambridge** | | | | **Chicago** | | | |
|---|---|---|---|---|---|---|---|---|
| | **Mean** | **Median** | **SD** | **Range** | **Mean** | **Median** | **SD** | **Range** |
| Age | 32.2 | 31.0 | 9.1 | 18 - 49 | 29.6 | 26.0 | 8.5 | 19 - 43 |
| NART IQ | 109.6 | 110.28 | 6.8 | 87 - 119 | 104.4 | 104.8 | 5.7 | 94 - 118 |
| BIS-11 [a] | 63.9 | 63.5 | 9.5 | 48 - 85 | 55.8 | 51.0 | 11.0 | 40 - 75 |
| Padua Inventory [a] | 10.7 | 8.5 | 9.3 | 0 - 32 | 9.5 | 7.0 | 7.2 | 1 - 25 |
| Gender (M:F) | 18 : 4 | | | | 8 : 5 | | | |
| **Education Level** | **n** | **%** | | | **n** | **%** | | |
| Some College | 7 | 31.8 | | | 5 | 38.5 | | |
| College degree | 6 | 27.3 | | | 3 | 23.1 | | |
| Post-College | 9 | 40.9 | | | 3 | 23.1 | | |

| | | | |
|---|---|---|---|
| Missing | / | 2 | 15.4 |

### Supplemental data analyses

#### Data Quality Control

Data was screened for quality, and subjects excluded according to the following criteria: i) Completely deterministic responding, i.e., if participants always gave the same response to each cue (e.g. always Go for Win cues and NoGo for avoid cues), indicating failure to understand the task. No subject met this criterion. ii) More than 25% (i.e., >40 trials) of pressing an uninstructed button, given that on those trials participants did not receive feedback and could not learn. This led to exclusion of one participant with 50 false responses resulting in a final sample size of *N* = 35. After exclusion of this one participant, the mean number of false responses was < 1 % across participants. Those 82 trials were subsequently excluded for the mixed model analysis (0.68% out of 12000 in total). For the logistic regression, we excluded trials on which participants showed reaction times (RT) faster than 200 ms, as it can be assumed that such fast RTs cannot reflect the stimulus being adequately processed (24 trials in total).

As noted in the main manuscript, we detected difference across test site in terms of performance, which are further visualized in Figure S1. Here, we also report the employed model equation for completeness:

Choice~ Req Action* Valence * Drug * Site + (Req Action* Valence * Drug * Site + 1|Subject)

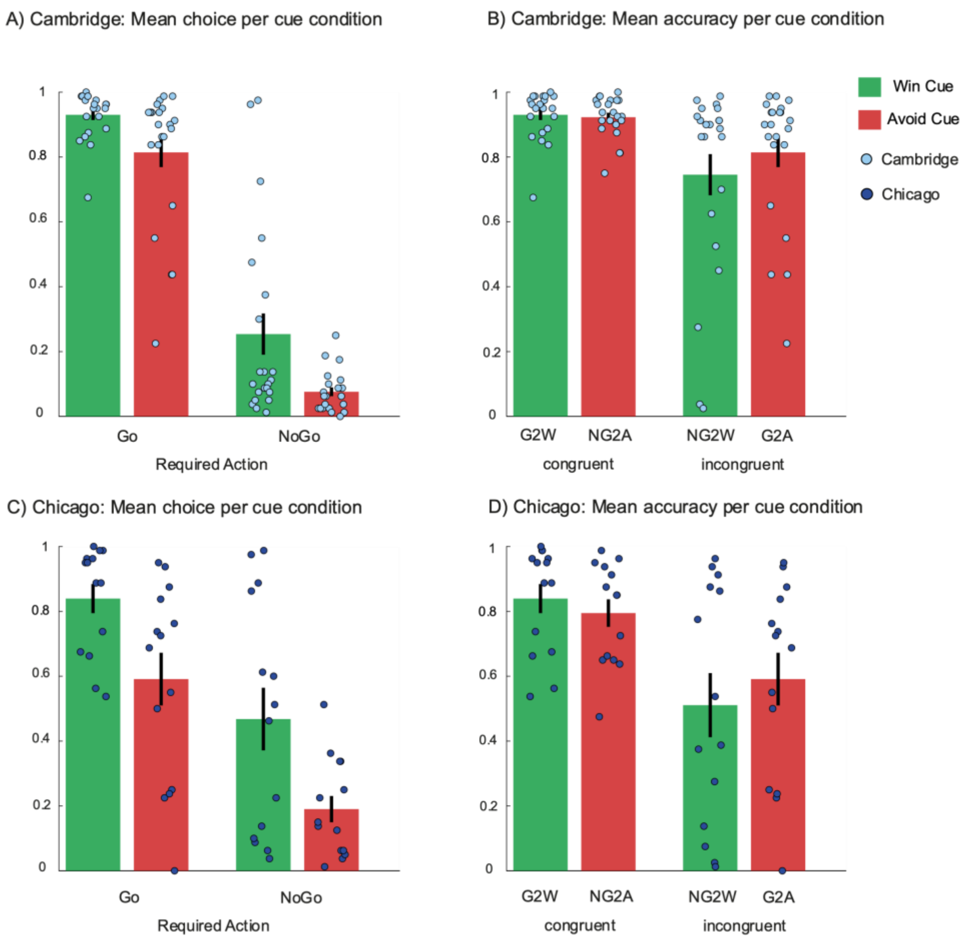

**Figure S1.** Site differences in performance. **A** and **B**) Mean probability of making a Go response (P(Go) for each cue condition and mean Accuracy per cue sorted according to bias congruent and incongruent response requirements, depicted for the Cambridge recruitment site. C and D) Mean probability of making a Go response (P(Go) for each cue condition and mean Accuracy per cue sorted according to bias congruent and incongruent response requirements, depicted for the Chicago recruitment site

#### Task block effects

In the main manuscript, we assessed whether the impact of tolcapone on motivational biases was constant over time. For this, we included task block (2 blocks of 80 trials) as 2-level factor, within-subject effect in the mixed model. The Valence x Drug x Block interaction was significant ($\chi^2(1) = 4.6$, p-value = 0.03). For completeness, we report the model equation and the full statistics of this model below. As reported in the main manuscript, we additionally analyzed the Drug x Valence interaction for block 1 and block 2 separately.

Choice~ Req Action* Valence * Drug * Block * Site + (Req Action* Valence * Drug * Block + 1|Subject)

**Table S1: Effects of task block**

| | ß estimates | SE | $\chi^2$ | p-value |
|---|---|---|---|---|
| **Main effects** | | | | |
| valence | -0.872 | 0.18 | 24.7 | <.001 *** |
| required action | 2.233 | 0.24 | 85.8 | <.001 *** |
| drug | -0.075 | 0.10 | 0.6 | 0.4 |
| site | -0.006 | 0.14 | <0.01 | 1 |
| block | 0.191 | 0.09 | 4.8 | .03* |
| **Interaction effects** | | | | |
| required action x valence | 0.013 | 0.08 | 0.02 | 0.9 |
| drug x site | 0.073 | 0.10 | 0.6 | 0.5 |
| drug x block | 0.148 | 0.07 | 4.0 | 0.05* |
| site x block | -0.110 | 0.09 | 1.6 | 0.2 |
| valence x drug | -0.174 | 0.10 | 3.1 | 0.1 |
| valence x site | 0.201 | 0.18 | 1.3 | 0.3 |
| valence x block | -0.009 | 0.07 | 0.02 | 0.9 |
| required action x drug | 0.151 | 0.13 | 1.3 | 0.3 |
| required action x site | 0.638 | 0.24 | 7.0 | 0.008 ** |
| required action x block | -0.385 | 0.10 | 14.6 | <.001 *** |
| drug x site x block | -0.136 | 0.07 | 3.8 | 0.1 |
| valence x drug x site | 0.094 | 0.10 | 0.9 | 0.3 |
| valence x drug x block | -0.114 | 0.05 | 4.6 | 0.03 * |
| valence x site x block | -0.078 | 0.07 | 1.2 | 0.3 |
| required action x drug x site | -0.085 | 0.13 | 0.4 | 0.5 |
| required action x drug x block | -0.027 | 0.07 | 0.2 | 0.7 |
| required action x site x block | -0.186 | 0.10 | 3.4 | 0.1 |
| valence x required action x drug | 0.026 | 0.07 | 0.1 | 0.7 |
| valence x required action x site | 0.078 | 0.08 | 0.9 | 0.4 |
| valence x required action x block | -0.167 | 0.07 | 5.8 | 0.02* |
| valence x drug x site x block | 0.056 | 0.05 | 1.1 | 0.3 |
| required action x drug x site x block | -0.104 | 0.07 | 2.3 | 0.1 |
| valence x required action x drug x site | 0.051 | 0.07 | 0.6 | 0.5 |
| valence x required action x drug x block | -0.040 | 0.06 | 0.4 | 0.5 |
| valence x required action x site x block | -0.087 | 0.07 | 1.6 | 0.2 |
| valence x required action x drug x block x site | -0.015 | 0.06 | <0.1 | 0.8 |

#### Session order effects

As a control analysis, we assessed whether session order (i.e. whether tolcapone was administered on testing day 1 or 2) interacted with the Drug x Valence interaction of interest. This was done given such effects were observed in a previous pharmacological study with the same task [6]. However, we did not see a Drug x Valence x Order interaction here ($\chi^2(1) = 1.05$, p-value = .3). Importantly, adding testing order did also not alter the significance of the Drug x Valence interaction ($\chi^2(1) = 6.2$, p-value = .01). For completeness, we report the full statistics of this analysis below (Table S2)

Choice~ Req Action* Valence *Order*Site+ (Req Action* Valence * Order + 1|Subject)

**Table S2: main and interaction effects of mixed model for choice data including drug, order and site.**

| | ß estimate | SE | $\chi^2$ | p-value |
|---|---|---|---|---|
| **Main effects** | | | | |
| valence | -0.814 | 0.15 | 31.7 | <.001 *** |
| required action | 1.955 | 0.20 | 100.4 | <.001 *** |
| drug | -0.086 | 0.08 | 1.3 | 0.3 |
| order | -0.347 | 0.10 | 11.2 | <.001 *** |
| site | -0.031 | 0.10 | <0.1 | 0.9 |
| **Interaction effects** | | | | |
| required action x valence | -0.006 | 0.07 | < 0.1 | 0.9 |
| drug x order | 0.058 | 0.08 | 0.6 | 0.5 |
| drug x site | 0.068 | 0.08 | 0.8 | 0.4 |
| order x site | 0.230 | 0.10 | 4.9 | 0.03 * |
| valence x drug | -0.193 | 0.08 | 6.2 | 0.01 * |
| valence x order | 0.292 | 0.15 | 4.1 | 0.04 * |
| valence x site | 0.161 | 0.15 | 1.2 | 0.3 |
| required action x drug | 0.098 | 0.11 | 0.7 | 0.4 |
| required action x order | 0.283 | 0.20 | 2.1 | 0.2 |
| required action x site | 0.594 | 0.20 | 9.3 | 0.002 ** |
| drug x order x site | 0.073 | 0.08 | 0.9 | 0.3 |
| valence x drug x order | -0.080 | 0.08 | 1.1 | 0.3 |
| valence x drug x site | 0.070 | 0.08 | 0.8 | 0.4 |
| valence x order x site | -0.177 | 0.15 | 1.5 | 0.2 |
| required action x drug x order | -0.157 | 0.11 | 1.9 | 0.2 |
| required action x drug x site | -0.117 | 0.11 | 1.1 | 0.3 |
| required action x order x site | -0.101 | 0.20 | 0.3 | 0.6 |
| valence x required action x drug | 0.035 | 0.06 | 0.3 | 0.6 |
| valence x required action x order | -0.095 | 0.07 | 1.9 | 0.2 |
| valence x required action x site | 0.030 | 0.07 | 0.2 | 0.7 |
| valence x drug x order x site | 0.111 | 0.08 | 2.0 | 0.2 |
| required action x drug x order x site | -0.032 | 0.11 | 0.1 | 0.8 |
| valence x required action x drug x order | 0.044 | 0.06 | 0.5 | 0.5 |
| valence x required action x drug x site | 0.004 | 0.06 | <0.1 | 1 |
| valence x required action x order x site | -0.001 | 0.07 | <0.1 | 1 |
| valence x required action x drug x order x site | 0.033 | 0.06 | 0.3 | 0.6 |

#### Mixed regression analysis of reaction times

As a secondary analysis, we analysed reaction times using the same factors (Required action, Valence, Drug as within subject variables, Site as between-subject variable), using linear mixed-effects models as implemented in lme4, which are appropriate for continuous RT data.

Note that this analysis has much lower power than the analysis of the choice data, as by definition we can only analyse Go responses.

RT ~ Req Action* Valence * Drug * Site + (Req Action* Valence * Drug + 1|Subject)

Results from this analysis conceptually replicate the main findings from the choice data regarding the non-drug effects. People were faster to respond to a Win rather than an Avoid cue (Valence $\chi^2(1) = 42.0$, p-value < .001), indicating a presence of a motivational bias in invigorating of responding. People were also faster to make a Go response when this was correct (i.e. for a Go cue), relative to when this was incorrect (i.e. for NoGo cue) (required action: $\chi^2(1) = 37.8$, p-value < .001), indicating that there was (in- or explicit) awareness of the correctness of making a Go response, i.e. learning. However, there were no interactions with tolcapone (Drug x Valence: $\chi^2(1) < 0.05$, p-value = .08). Full statistics are presented in Table S3.

**Table S3: main and interaction effects of mixed model for RT data**

| | ß estimates | SE | $\chi^2$ | p-value |
|---|---|---|---|---|
| **Main effects** | | | | |
| required action | -0.077 | 0.01 | 37.8 | <0.001*** |
| valence | 0.054 | 0.01 | 42.0 | < 0.001 *** |
| drug | -0.014 | 0.01 | 2.8 | 0.1 |
| site | 0.030 | 0.02 | 1.6 | 0.2 |
| **Interaction effects** | | | | |
| required action x valence | -0.011 | 0.01 | 1.3 | 0.3 |
| drug x site | -0.009 | 0.01 | 1.1 | 0.3 |
| valence x drug | -0.002 | 0.01 | < .05 | 0.8 |
| valence x site | 0.002 | 0.01 | 0.1 | 0.8 |
| required action x drug | 0.005 | 0.01 | 0.5 | 0.5 |
| required action x site | -0.017 | 0.01 | 1.8 | 0.1 |
| valence x drug x site | -0.012 | 0.01 | 3.3 | 0.07 + |
| required action x drug x site | 0.013 | 0.01 | 3.3 | 0.1 |
| valence x required action x drug | 0.006 | 0.01 | 0.9 | 0.3 |
| valence x required action x site | -0.005 | 0.01 | 0.5 | 0.5 |
| valence x required action x drug x site | 0.008 | 0.01 | 1.7 | 0.2 |

### Computational modelling analysis

#### Model equations

Across all models, action weights (w) for Go and NoGo responses were determined per trial (t) and cue (s) and used together with a softmax function to compute the associated choice probabilities:

$$p(a_t|s_t) = \frac{\exp\,(w(a_t, s_t))}{\sum_{a'} \exp\,(w(a', s_t))} \qquad (1)$$

The model M1 was a standard Rescorla Wagner model that only included two model parameters, namely a learning rate (ε) and a feedback sensitivity parameter (ρ). Action weight for both motor responses were derived from unbiased action values using a standard delta-learning rule including these two parameters.

$$Q_t(a_t, s_t) = Q_{t-1}(a_t, s_t) + \varepsilon(\rho r_t - Q_{t-1}(a_t, s_t)) \qquad (2)$$

For the second model (M2), an additional parameter capturing the 'go' bias (b) was introduced. The go bias parameter *b* gets added exclusively to the action weight of the Go response and describes an individuals' tendency to make a 'go' response, independent of cue valence:

$$w_t(a,s) = \begin{cases} Q_t(a,s) + b & if\ a = Go \\ Q_t(a,s) & else \end{cases} \tag{3}$$

To also assess the latent mechanisms in the decision process during tolcapone and placebo administration, we additionally employed computational modelling. For this, Model M3 was extended by a bias parameter π which was introduced to capture the impact of motivational biases on decision making. Modelling the impact of motivational biases was achieved by allowing the anticipated cue value (V) to impact the action weights, either by increasing or decreasing the action weight of the Go responses. Given that in this task, the cue valence for Win and Avoid cues was cued, we fixed the weight of the Go response to be positive (V = + 0.5) for Win cues or negative for Avoid cues (V = -0.5). Thereby, the strength of the motivational bias influencing action selection can be derived from the bias parameter π:

$$w_t(a,s) = \begin{cases} Q_t(a,s) + b\ +\ V\pi(s) & if\ a = Go \\ Q_t(a,s) & else \end{cases} \tag{4}$$

Except for the learning rates ε, which were constrained between 0 and 1 (via a probit transform), all other parameters were left unconstrained.

#### Model Fitting and Comparison

Model fitting was performed with the cbm toolbox implemented in matlab [7], employing Hierarchical Bayesian Inference (HBI). Hierarchical Bayesian Inference (HBI) treats model as a random rather than a fixed effects term [8,9]. Thus, individual data can impact updating of group-level parameters differently, with participants whose observed data is fitted better by the respective model impacting the update to a greater extent. To determine the best fitting model, we employed a random effects model comparison [8,9] based on the Laplace approximation of model evidence on the individual level[10,11] from which group evidence is subsequently computed. We first compared model evidence across base models M1-M3 using model frequency and protected exceedance probability (PXP) [7]. PXP compares which model is expressed most often [8] while controlling for the possibility that this finding could result from chance. Simulations confirmed that the winning model could qualitatively reproduce key patterns of the data (see details below).

#### Model comparison and parameter estimates

We already report the modelling results in the main manuscript and illustrate the results from the winning tolcapone model M4 in Figure 2 in the main manuscript. For completeness, we here report all parameter estimates from the three base models (M1-M3) as well as the numerical results from the base and full model comparisons in Table S4.

| | **M1**<br>**Basic RL** | **M2**<br>**+ Go bias** | **M3**<br>**+ Pavlovian bias** | **M4**<br>**+ TOLC** |
|---|---|---|---|---|
| **Parameter estimates** | | | | |
| Sensitivity ρ | 8.58 [1.01] | 8.26 [1.09] | 7.91 [1.07] | pla: 8.06 [0.95] |

| | | | | |
|---|---|---|---|---|
| | | | | tol: 7.36 [1.07] |
| Learning rate ε | 0.25 [0.05] | 0.28 [0.05] | 0.29 [0.04] | pla: 0.23 [0.03]<br>tol: 0.28 [0.03] |
| Gobias b | | 0.19 [0.09] | 0.24 [0.11] | pla: 0.17 [0.14]<br>tol: 0.25 [0.11] |
| Pavlovian bias π | | | 1.32 [0.26] | pla: 1.80 [0.30]<br>tol: 1.19 [0.25] |
| **Base Model Comparison (M1 – M3, excluding tolcapone model M4)** | | | | |
| Model frequency (%) | 25.3 | 31.8 | 42.9 | |
| PXP | 0.07 | 0.21 | 0.72 | |
| **Full Model Comparison (M1 – M4)** | | | | |
| Model frequency (%) | 12.5 | 15.6 | 11.4 | 60.5 |
| PXP | <0.001 | <0.001 | <0.001 | 0.998 |

**Table S4:** Overview of transformed hierarchical model parameter estimates (M [SE]) and hierarchical model comparison metrics. Protected exceedance probability (PXP) indicated that the winning model for the base comparison (M1-M3) was model M3 comprising 4 parameters including rho, epsilon, go and a motivational bias parameter. Full model comparison including the model M4, allowing for a drug modulation impact on parameters, showed that this model outperformed the simpler model M3. PXP = protected exceedance probability. The learning rate was transformed using a probit transformation restricted it to values between 0 and 1.

#### Control analysis of site effect for estimated model parameters

In line with the analysis of the raw behavioural data, we also assessed whether site interacted with the tolcapone effect on the Pavlovian bias parameter but also the other three model parameters derived from the winning computational model using the following general model equation:

Parameter~ Drug * Site

Table S5 reports the full regression analysis results with factors Drug and Site, which are already referred to in the main manuscript. Apart from the already reported effect of tolcapone on the Pavlovian bias parameter ($\chi^2(1) = 5.4$, p-value = .02), we report significant effects of testing site on both learning rate ( $\chi^2$ (1)= 8.1, p = .005)and feedback sensitivity ( $\chi^2$ (1)= 11.3, p < .001) which can capture the difference in performance in terms of instrumental learning of the task, but are unrelated to the effects of tolcapone. For further full results, see table S6. Effects are visualized in Figure S2.

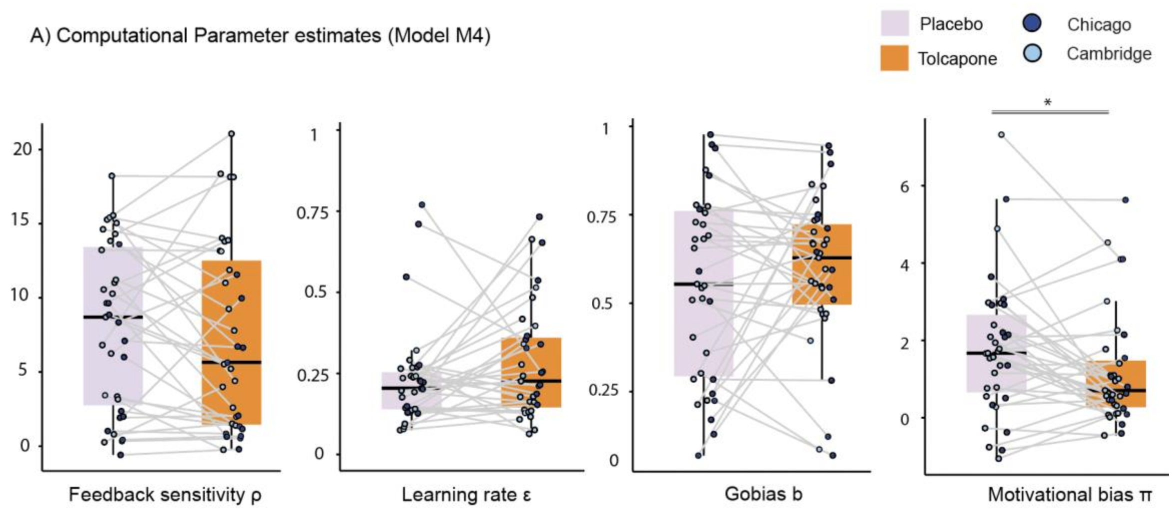

**Figure S2:** A) Tolcapone effect on parameter estimates for motivational bias parameter for the placebo and tolcapone session as well as the parameters for feedback sensitivity, the learning rate and the Go bias from the winning model M4.

**Table S5 Drug effects on model parameter estimates for observed data**

| parameter | | $\chi^2$ | **p-value** |
|---|---|---|---|
| sensitivity ρ | ***Main effects*** | | |
| | drug | 0.8 | .4 |
| | site | 11.3 | <.001 *** |
| | ***Interaction effects*** | | |
| | drug x site | 0.1 | .8 |
| learning rate ε | **Main effects** | | |
| | drug | 1.8 | .2 |
| | site | 8.1 | .005 ** |
| | **Interaction effects** | | |
| | drug x site | 0.04 | .8 |
| Go bias b | **Main effect** | | |
| | drug | 1.1 | .3 |
| | site | 0.1 | .7 |
| | **Interaction effect** | | |
| | drug x site | 0.4 | .5 |
| Pavlovian bias π | **Main effect** | | |
| | drug | 5.4 | .02 * |
| | site | 0.2 | .6 |
| | **Interaction effect** | | |
| | drug x site | < 0.1 | .9 |

#### Model validation and parameter recovery

While model comparison can indicate which model performs best within a selected model-space, it does not answer whether a model is able to reproduce key qualitative patterns in the data. [12]. Thus, we also tested whether our model could recover the model parameters. For this, we simulated 150 synthetic datasets per individual, based on their individual optimal parameter estimates from the winning model M4. We first validated that this simulated data from the winning model captures the key features of the task. As can be observed in Figure S3A, the model captured both instrumental learning and the presence of a motivational bias (green above red lines), and matched the original data relatively well.

We then re-fitted our winning computational model to each of the 150 simulated datasets and averaged the resulting model parameter estimates across simulation runs, and examined the difference in parameter estimates for the Tolcapone and Placebo condition. This recovered the significant difference in the Pavlovian bias parameter π ($\chi2(1) = 6.3$, p-value = .01; Fig S2). The feedback sensitivity and go bias parameters did not differ between placebo and tolcapone (for observed and simulated data, all $p > .1$). However, there was a significant difference between the recovered learning rate under tolcapone versus placebo ($\chi2(1) = 5.0$, p-value = .03), which was not significant when fitting the real data ($\chi2(1) = 1.8$, p=0.2). However, for both real and simulated data, the estimated learning rate decreased under tolcapone. It is possible that the non-significant parameter difference for the real data surfaced as a significant effect in the simulated data. This is because model fitting 'reduces' the real data to a few parameters, effectively getting rid of any variance not explained by the model. Thus, simulations result in 'cleaner', i.e. more consistent, data. This can then lead to parameter estimates to become significant. Speculatively, this change in the instrumental learning rate (in addition to a modulation of motivational bias parameter) is putatively of interest, in that it is in line with a shift in the balance between the instrumental and Pavlovian controllers.

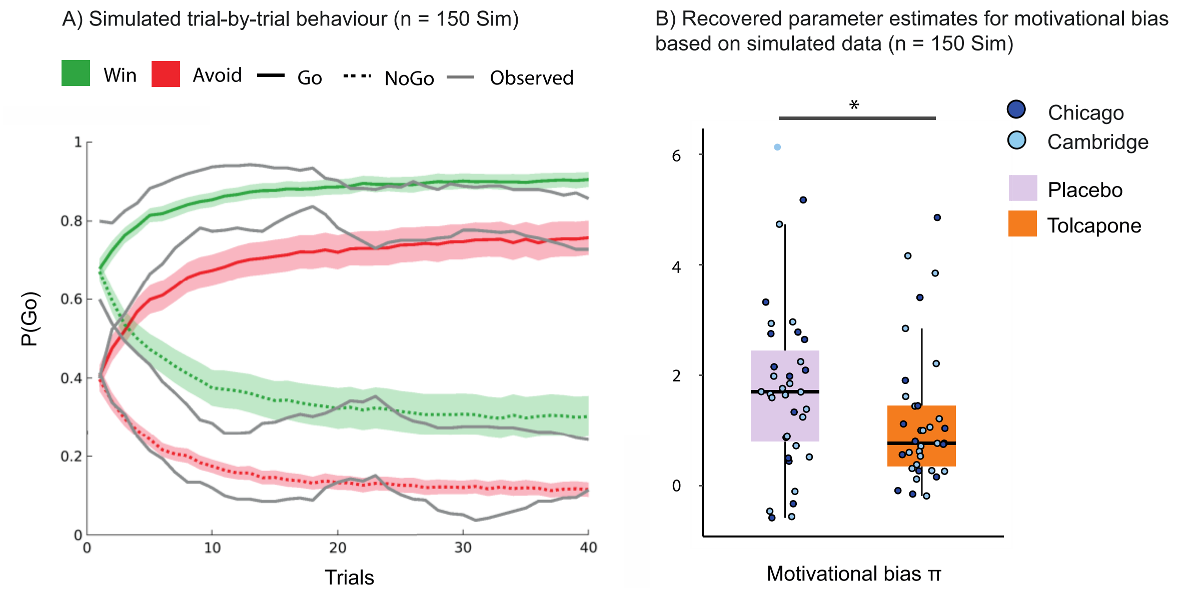

**Figure S3:** A) Simulated trial by trial behaviour based on n = 150 simulated datasets. B) Recovered parameter estimate for motivational bias parameter for the placebo and tolcapone session based on n = 150 simulation runs.
